# Elucidation of Global Trends in the Effects of VX-661 and VX-445 on the Expression of Clinical CFTR Variants

**DOI:** 10.1101/2022.10.14.512300

**Authors:** Andrew G. McKee, Eli F. McDonald, Wesley D. Penn, Charles P. Kuntz, Karen Noguera, Laura M. Chamness, Francis J. Roushar, Jens Meiler, Kathryn E. Oliver, Lars Plate, Jonathan P. Schlebach

**Affiliations:** Department of Chemistry, Indiana University, Bloomington, Indiana, USA; Department of Chemistry, Vanderbilt University, Nashville, TN, USA; Center for Structural Biology, Vanderbilt University, Nashville, TN, USA; Institute for Drug Development, Leipzig University, Leipzig, SAC, Germany; Department of Pediatrics, Emory University School of Medicine, Atlanta, GA, USA; Department of Biological Sciences, Vanderbilt University, Nashville, TN, USA

**Keywords:** Cystic Fibrosis, CFTR, Corrector, Membrane Protein Folding, Misfolding, Proteostasis

## Abstract

Cystic fibrosis (CF) is a chronic genetic disease caused by mutations that compromise the expression and/ or function of the cystic fibrosis transmembrane conductance regulator chloride channel (CFTR). Most people with CF harbor a common misfolded CFTR variant (ΔF508), which can be rescued by combination therapies containing “corrector” compounds that restore its expression. Nevertheless, there are over 400 other CF variants that differ in their sensitivity to correctors for reasons that remain unclear. In this work, we utilize deep mutational scanning to quantitatively compare the effects of two FDA-approved correctors on the plasma membrane expression of 129 known CF variants, including 45 that are currently unclassified. Across 67 variants with attenuated expression, we find that VX-661-sensitive variants generally exhibit intermediate expression and feature mutations near its binding pocket in transmembrane domains (TMDs) 1, 2, 3, and 6. VX-445 also primarily rescues variants with intermediate expression but is instead uniquely effective towards mutations near its binding pocket in TMDs 10 & 11. Structural calculations suggest corrector binding provides similar stabilization to both sensitive and insensitive variants. These findings collectively suggest the mutation-specific effects of these compounds depend on the degree of variant destabilization and/ or the timing of cotranslational folding defects. Combining these correctors synergistically rescues variants with deficient and intermediate expression alike, presumably by doubling the total binding energy and suppressing defects at different stages of translation. These results provide an unprecedented overview of the properties of rare CFTR variants and establish new tools for CF pharmacology.

## Introduction

Cystic fibrosis (CF) is a common genetic disease caused by an array of loss-of-function mutations in the cystic fibrosis transmembrane conductance regulator gene (*CFTR*), which encodes the CFTR chloride channel protein. Over 400 CFTR mutations have been definitively linked to CF to date (https://www.cftr2.org/). Nevertheless, approximately 85% of CF patients carry at least one copy of the ΔF508 variant, which both reduces the expression of the channel and compromises its functional gating.^1^ The remaining ∼15% of CF patients bear combinations of rare CF mutations that can attenuate CFTR function by either compromising the CFTR transcript (class I), enhancing CFTR misfolding and degradation in the endoplasmic reticulum (class II), disrupting channel gating (class III & IV), reducing the total accumulation of mature CFTR (class V), reducing its stability at the plasma membrane (class VI), or triggering some combination of these molecular defects.^2^ Though many of the known CF variants remain uncharacterized, current evidence suggests the majority of rare CF variants generate class II folding defects.^2^ For this reason, efforts to develop CF therapeutics have focused extensively on approaches to rescue the folding and expression of destabilized CFTR variants.^3,4^

A general understanding of the molecular effects of CF mutations has facilitated the recent development and clinical targeting of corrector and potentiator molecules that enhance CFTR expression and function, respectively.^5^ Clinically approved type I correctors such as VX-809 (Lumacaftor®) and VX-661(Tezacaftor®) bind within the first membrane spanning domain (MSD1) and suppress cotranslational misfolding, which ultimately enhances the plasma membrane expression (PME) of a certain misfolded CF variants.^6,7^ These compounds are most effective at restoring the function of misfolded variants when combined with potentiators such as VX-770 (Ivacaftor®),^8,9^ which binds to CFTR’s eighth transmembrane domain (TMD) and enhances the opening of the channel.^10^ A more recently approved CFTR modulator known as VX-445 (Elexacaftor®), which binds within TMDs 10 & 11 and acts as both a corrector and potentiator,^11-13^ provides a complementary benefit to previously developed modulators. The current leading therapeutic (Trikafta®), which consists of a combination of VX-661, VX-445, and VX-770, can be used to treat most people with CF.^9^ However, this treatment is not universally tolerated in the clinic and is not effective against certain *CFTR* mutations.^14^ For example, two of the five most common CF mutations (N1303K and ΔI507) do not appreciably respond to these molecules for reasons that remain unclear. Fortunately, a variety of new CFTR modulators are advancing through clinical trials. These modulators could potentially expand the matrix of combinatorial therapies and address certain unmet medical needs. Nevertheless, identifying the optimal combination therapies for individual variants will become increasingly challenging given the limited throughput of the current gold-standard western blot and electrophysiology-based approaches that are most often used to survey the effects of CFTR modulators on CF variant expression and or function, respectively. New approches to efficieciently compare the mutation-specific effects of CFTR modulators are thus needed to efficiently profile the response of rare CF variants to combinations of small molecules.

In this work, we use deep mutational scanning (DMS) to compare the effects of VX-661, VX-445, and a combination of both molecules on the PME of 129 of the most common CF variants, including 53 without previously documented experimental characterization. We find that 67 of 105 potentially correctable missense and in-frame insertion/ deletion (indel) variants in this library exhibit diminished PME relative to wild-type (WT) CFTR in HEK293T cells. Our results reveal that, on average, mutations within the first membrane-spanning domain (MSD1) or the first nucleotide binding domain (NBD1) cause the greatest reduction in PME. PME measurements in the presence of VX-661 or VX-445 reveal that these compounds are each most effective towards variants with intermediate expression that perturb regions near their respective binding pockets. However, combining these molecules rescues mutants with varied expression throughout the CFTR structure. The overall pharmacological profiles of these variants suggest rescue primarily depends on residual PME. Nevertheless, we note that the regiospecific effects of VX-661 and VX-445 is consistent with recent evidence suggesting that their synergistic interaction arises from their differential stabilization of early and late-stage cotranslational folding intermediates, respectively.^13^ Our findings provide a broad overview of the properties of CF variants and provide insights into the molecular effects of correctors.

## Results

### Plasma Membrane Expression of Rare Cystic Fibrosis Variants

To profile the PME of CF variants by DMS, we first generated a one-pot library of mutants bearing a previously described triple-HA tag in ECL4 that enables robust CFTR immunostaining at the plasma membrane.^4^ We generated a series of CFTR expression vectors encoding individual mutants from the CFTR2 database (https://www.cftr2.org/) in combination with a unique molecular identifier (UMI) sequence in the plasmid backbone, which we used to track each variant with Illumina sequencing in the downstream assay. The final library consists of a stoichiometric pool of 129 vectors that collectively encode 104 missense mutations, 16 nonsense mutations, 6 synonymous mutations, and 3 indel mutations (Table S1). This collection of variants includes 14 known class I mutations, 46 class II mutations, 14 class III mutations, and 45 mutations with previously undocumented classifications (Table S1). Our manual assembly of this library provides opportunities to incorporate additional variants of interest in the future. Additionally, the equivalent abundance of each mutant in the pooled library ensures even sampling of each variant in the downstream assay, which ultimately improves the precision of the resulting PME measurements.

To survey the effects of these mutations on CFTR PME, we measured their surface immunostaining in parallel by DMS as previously described.^15-17^ Briefly, we first used our genetic library to generate a pool of recombinant HEK293T cells that each inducibly express a single CFTR variant from a defined genomic locus. The surface immunostaining profile of these recombinant cells reveals that ∼30% express variants with comparable immunostaining to WT CFTR (Int. = 1695 ± 291, Table S1) while the remaining ∼70% express variants that exhibit immunostaining intensities similar to the misfolded ΔF508 variant (Int. = 188 ± 28, Fig. 1A). To estimate the PME of individual CF variants, we utilized fluorescence activated cell sorting (FACS) to fractionate this mixed cell line based on the immunostaining intensity of their expressed CFTR variants. We then extracted the genomic DNA from each cellular fraction and used Illumina sequencing to quantify the relative abundance of the UMI’s associated with each variant in each fraction. Sequencing data were then used to infer the relative surface immunostaining intensities of each variant as previously described.^16^

**Figure 1.**
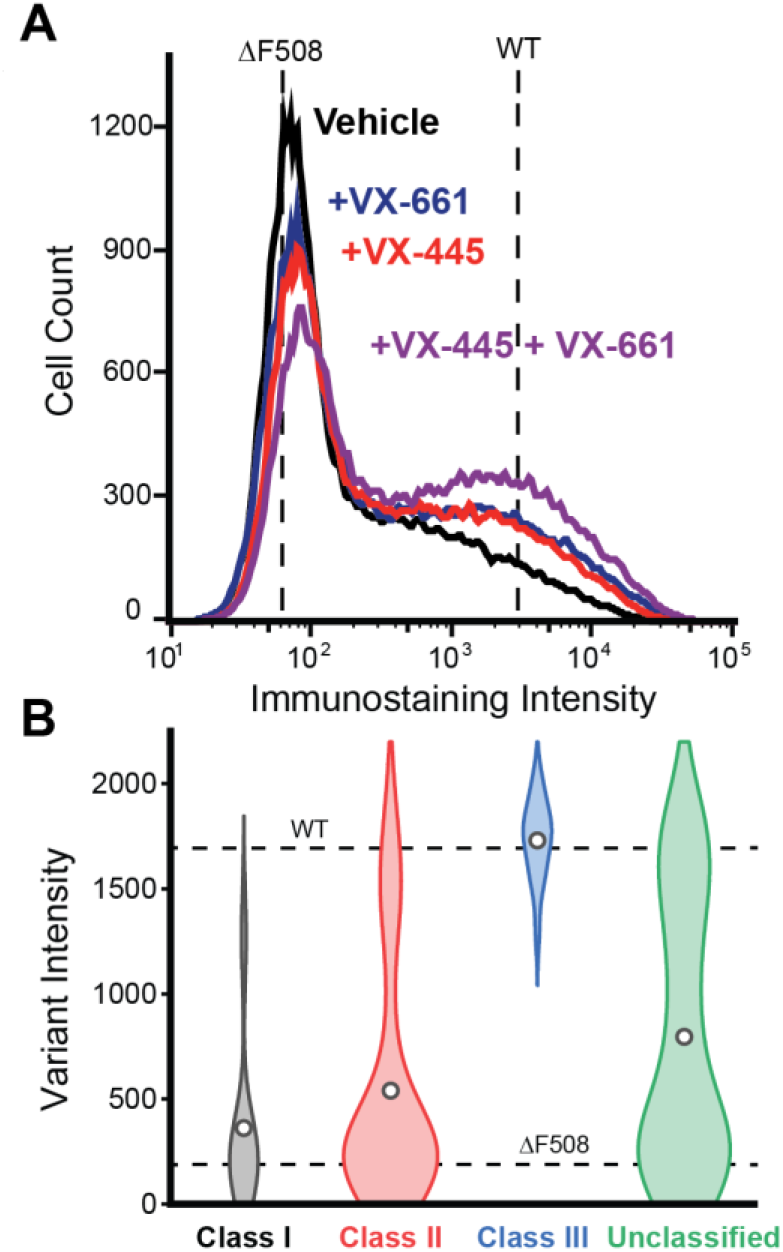
Surface Immunostaining Profiles of Recombinant Cells Expressing Known CF Variants. A) A histogram depicts the distribution of CFTR surface immunostaining intensities for recombinant HEK293T cells collectively expressing a library of 129 CFTR variants dosed with either vehicle (black), 3 μM VX-661 (blue), 3 μM VX-445 (red), or 3μM of both compounds (purple). B) A violin plot depicts the distribution of immunostaining intensities among known class I mutants (n=14, gray), class II mutants (n=46, red), class III mutants (n=14, blue), and unclassified (n=45, green) mutants. Open circles indicate the average intensity values. The mean immunostaining intensity of recombinant cells expressing WT or ΔF508 CFTR are shown with dashed lines in each panel for reference.

As expected, individual surface immunostaining intensities are correlated with the intensity of the mature glycoform levels of select variants (Pearson’s *R* = 0.61, *p* = 3.8 × 10^−5^, Figure S1). Additionally, the six synonymous variants in our library have immunostaining intensities within 4% of the WT value (Table S1), which confirms these mutations have minimal impact on CFTR PME. The precision of these measurements is also demonstrated by the consistent immunostaining intensities of three clinical missense variants that each generate an S549R substitution (72-74% of the WT intensity, Table S1). Interestingly, we find that only 12 of 16 nonsense mutations exhibit little to no immunostaining. The remaining four fall within NBD2 and exhibit marked PME (Table S1), which is consistent with previous observations suggesting CFTR variants bearing certain NBD2 truncations can still traffic to the plasma membrane.^18,19^ Consistent with expectations, most class II variants exhibit decreased surface immunostaining while the surface immunostaining intensities of known class III-only variants are generally comparable to WT (Fig. 1B). Previously unclassified variants diverge considerably with respect to their immunostaining intensity, which indicates that these mutations cause a spectrum of molecular effects (Fig. 1B). Together, these results demonstrate that DMS measurements can differentiate the molecular effects of CF mutations with high precision, and that the resulting measurements are generally consistent with previous experimental classifications.

Our library contains 105 potentially correctable missense and indel variants that vary considerably with respect to their surface immunostaining (Fig. 2A). Mutations within the lasso domain, MSD1, or NBD1 generally compromise PME more, on average, than those within the R-domain, MSD2, or NBD2 (Fig. 2B), which reflects the importance of early cotranslational assembly events in CFTR proteostasis.^20-23^ Indeed, a projection of variant intensity values onto the CFTR structure shows that many poorly expressed variants feature mutations within MSD1 or NBD1 (Fig. 2C). Nevertheless, the PME values for individual mutations within these domains vary widely, and there are both poorly expressed and properly expressed variants within each subdomain (Fig. 2D). Together, these observations echo general knowledge on the constraints of CFTR proteostasis while revealing tremendous heterogeneity in the quantitative effects of individual CF mutations. In the following we will focus our analysis on the sensitivity of these 105 variants to correctors.

**Figure 2.**
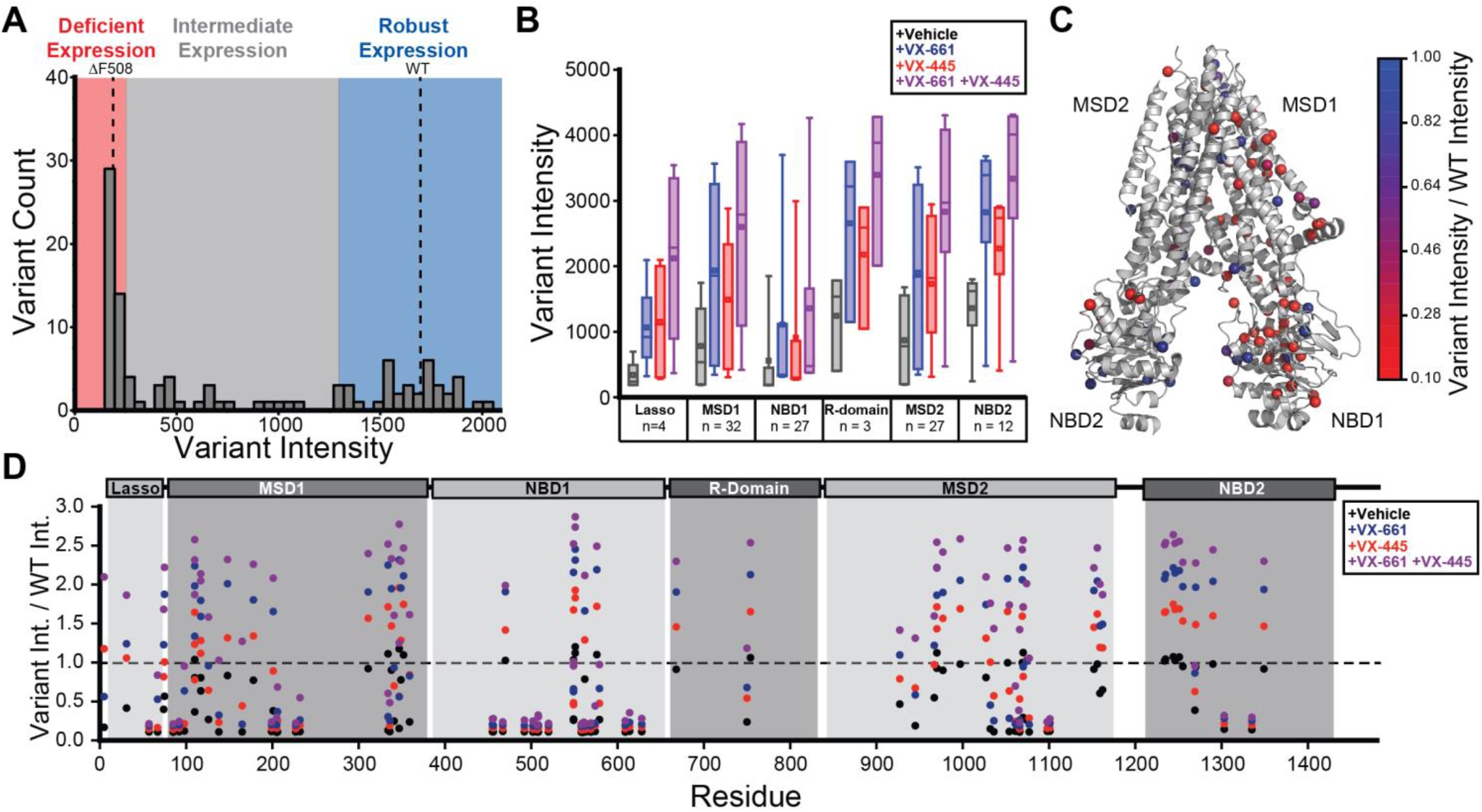
Surface Immunostaining Intensities of Individual CFTR Variants. The surface immunostaining intensities of individual variants were determined in by deep mutational scanning (DMS). A) A histogram depicts the distribution of surface immunostaining intensities across the 105 missense and indel variants in the presence of vehicle. Variants that exhibit “deficient,” “intermediate,” or “robust” plasma membrane expression fall within the red, gray, and blue regions, respectively. B) A box and whisker plot depicts the range of immunostaining intensities for variants within each subdomain in the presence of either vehicle (black), 3 μM VX-661 (blue), 3 μM VX-445 (red), or 3 μM of both compounds (purple). The upper and lower edges of the boxes represent the 75^th^ and 25^th^ percentile values, respectively. The upper and lower whiskers represent the 90^th^ and 10^th^ percentile values, respectively. The hash mark and square represent the median and mean values, respectively. C) Intensity values for individual variants in the presence of vehicle were normalized relative to the WT intensity and projected onto their mutated side chains in the CFTR closed state structure (PDB 5UAK). Side chain C_β_/ glycine H atoms are rendered as spheres and colored according to the average intensity from three replicate DMS experiments. D) Variant immunostaining intensities in the presence of vehicle (black), 3 μM VX-661 (blue), 3 μM VX-445 (red), or 3 μM of both compounds (purple) were normalized relative to the WT intensity in the presence of vehicle and plotted against their residue number. The positions of domain boundaries are shown for reference.

### Impact of VX-661 on CFTR Variant Plasma Membrane Expression

The response of rare CF variants to different corrector compounds varies considerably for reasons that remains poorly understood. To compare how these variants respond to a type I corrector, we repeated these experiments in the presence of VX-661. Incubating recombinant cells expressing CFTR variants with 3 μM VX-661 generally enhances the proportion of cells with WT-like surface immunostaining (Fig. 1A), though the effects of this compound on individual mutants deviates considerably (Fig. 2D). Rescue appears to be generally associated with variant PME (Figs. 3A & 4A). To evaluate the effects of VX-661 in relation to variant PME, we binned these variants according to their immunostaining as follows. We grouped 44 variants with deficient PME comparable to ΔF508 (Int. < 250), 38 variants with robust PME comparable to class III variants and WT (Int. > 1,300), and 23 variants with intermediate PME (250 < Int. < 1,300, Fig. 2A). Variants with intermediate PME are generally most sensitive (Figs. 3A & 4A)-all 23 exhibit gains equivalent to at least 25% of the WT immunostaining intensity. In contrast, only one in 44 variants with deficient PME reach this level of correction. This observation implies the degree of destabilization caused by variants with deficient PME exceeds the stabilization generated by VX-661 binding (see *Discussion*). Consistent with previous observations, VX-661 also generates measurable increases in the intensity of variants with robust PME as well, which potentially reflects the intrinsic instability of WT CFTR (Figs. 3A & 4A, see *Discussion*). Interestingly, we note that there are variants within each CFTR subdomain that exhibit measurable response to VX-661 (Fig. 2 B & D). However, a projection of fold-change intensity values onto the three-dimensional structure of CFTR reveals that VX-661 sensitive variants are most prevalent within MSD1 (Fig. 4B). Moreover, variants that exhibit the largest change in PME are clustered around the VX-661 binding pocket (Fig. 4C). These include Q98R and P67L-two variants that were previously found to be highly sensitive to type I correctors.^24,25^ This observation suggests VX-661 binding is most tightly coupled to the folding of the MSD1 subdomain, which is consistent with the findings of our recent molecular modeling efforts of VX-809-bound ΔF508 CFTR.^26^ Together, these findings reveal that VX-661 sensitivity is most pronounced among variants with intermediate PME bearing mutations near the VX-661 binding pocket.

**Figure 3.**
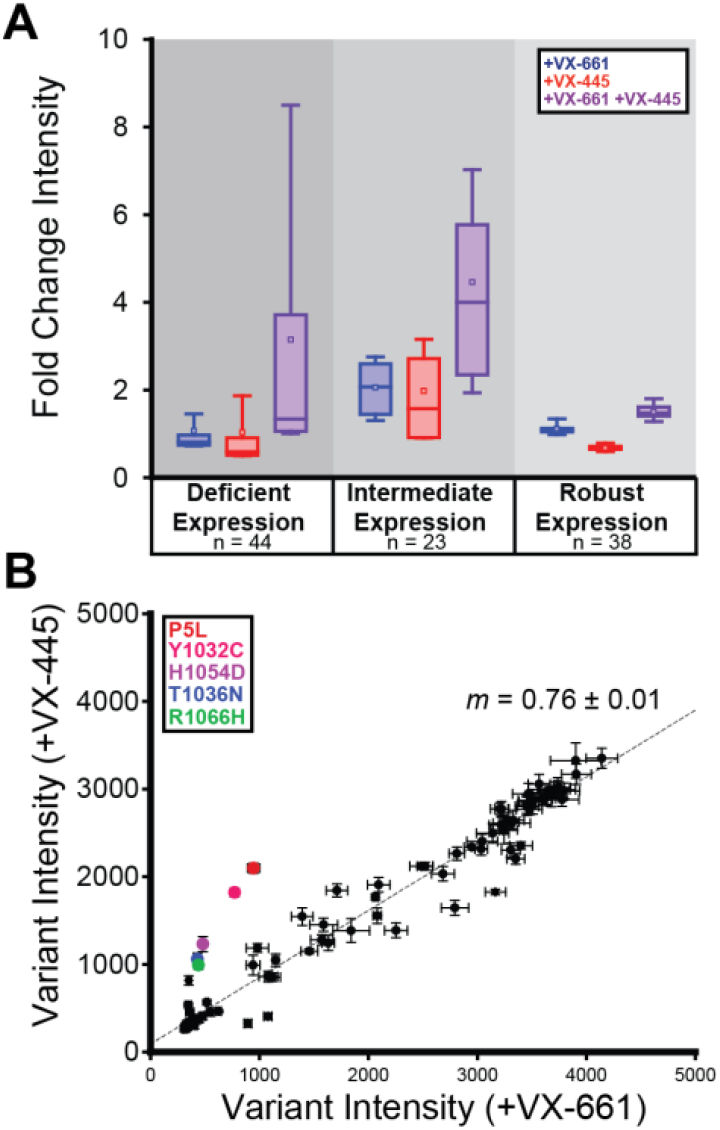
Comparison of the Effects of Correctors on the Plasma Membrane Expression of CFTR Variants. A) A box and whisker plot depicts the extent to which 3 μM VX-661 (blue), 3 μM VX-445 (red), or 3 μM of both compounds (purple) enhance the surface immunostaining intensities of variants with deficient expression (left), intermediate expression (center), and robust expression (right). The upper and lower edges of the boxes represent the 75^th^ and 25^th^ percentile values, respectively. The upper and lower whiskers represent the 90^th^ and 10^th^ percentile values, respectively. The hash mark and square represent the median and mean values, respectively. B) Variant immunostaining intensities in the presence of VX-445 are plotted against the corresponding intensities in the presence of VX-661. Measurements represent the average of three biological replicates and error bars reflect the standard deviation. A linear fit (gray dashes) and its fitted slope (*m*) are shown for reference.

**Figure 4.**
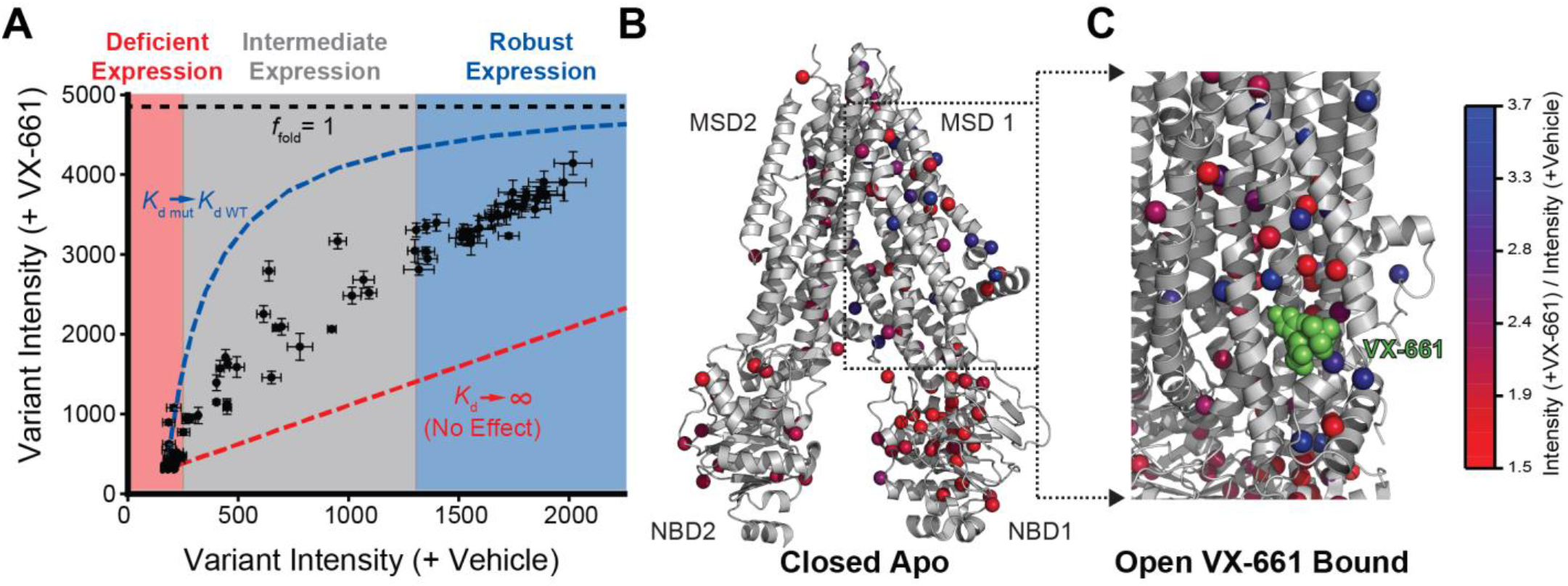
Impact of VX-661 on the Plasma Membrane Expression of CFTR Variants. Surface immunostaining intensities for individual variants were determined in the presence and absence of 3 μM VX-661 using deep mutational scanning (DMS). A) Variant immunostaining intensities in the presence of VX-661 are plotted against the corresponding intensities in the presence of vehicle. The relative positions of variants with deficient (red), intermediate (gray), and robust (blue) plasma membrane expression are shown for reference. Thermodynamic projections for the minimum (red dashes) and maximum shifts (blue dashes) in variant intensities are shown based on the projected surface immunostaining intensity of fully folded CFTR (*f*_fold_ = 1, black dashes). Measurements represent the average of three biological replicates and error bars reflect the standard deviation. B) The fold-change in intensity values for individual variants in the presence of VX-661 are projected onto their mutated side chains in the CFTR closed state structure (PDB 5UAK). Side chain C_β_/ glycine H atoms are rendered as spheres and colored according to the average fold-change from three biological replicates. C) The fold-change in intensity values for individual variants in the presence of VX-661 are projected onto a cutaway of the MSD1 region within the VX-661-bound CFTR active state structure (PDB 7SV7). The bound VX-661 is shown in green.

### Impact of VX-445 on CFTR Variant Plasma Membrane Expression

The current leading therapeutic (Trikafta ®) leverages VX-661 in combination with VX-445, a more recently developed CFTR modulator that synergistically enhances rescue through its effects as a potentiator and type III corrector.^11,12,14^ To gain insights into the basis for its complementary effects, we collected DMS measurements in the presence of VX-445. Incubating recombinant cells expressing CFTR variants with 3 μM VX-445 generates an increase in the proportion of cells with WT-like surface immunostaining (Fig. 1A). However, the effects of VX-445 are slightly muted relative to those of VX-661 (Fig. 3B). Nevertheless, we note that, like VX-661, VX-445 also fails to rescue most variants with deficient PME and is instead most effective towards variants with intermediate PME (Figs. 3A & 5A). This again suggests that, like VX-661, the response to VX-445 partially depends upon the underlying stability of each variant. As is true for VX-661, VX-445-sensitive variants are not confined to a single CFTR subdomain (Fig. 2 B & D). However, a projection of the fold-change intensity values onto the structure reveals that VX-445-sensitive mutations perturb distinct structural regions relative to VX-661-sensitive variants (compare Figs. 4B & 5B). Notably, we identify a cluster of poorly expressed variants in TMD10 (Y1032C, T1036N, and H1054D) and TMD11 (R1066H) that are relatively insensitive to VX-661 yet exhibit a five to seven-fold increase in PME in the presence of VX-445 (Fig. 3B). These mutations flank the VX-445 binding site and perturb two domain-swapped TMDs within MSD2 that form native contacts in MSD1 (Fig. 5C).^13^ VX-445 is also particularly potent towards the lasso domain variant P5L (Fig. 3B, Table S1), which is proximal to the VX-445 binding site (Fig. 5C).^13^ Thus, both VX-661 and VX-445 are most potent towards variants with intermediate expression bearing mutations near their respective binding pockets.

**Figure 5.**
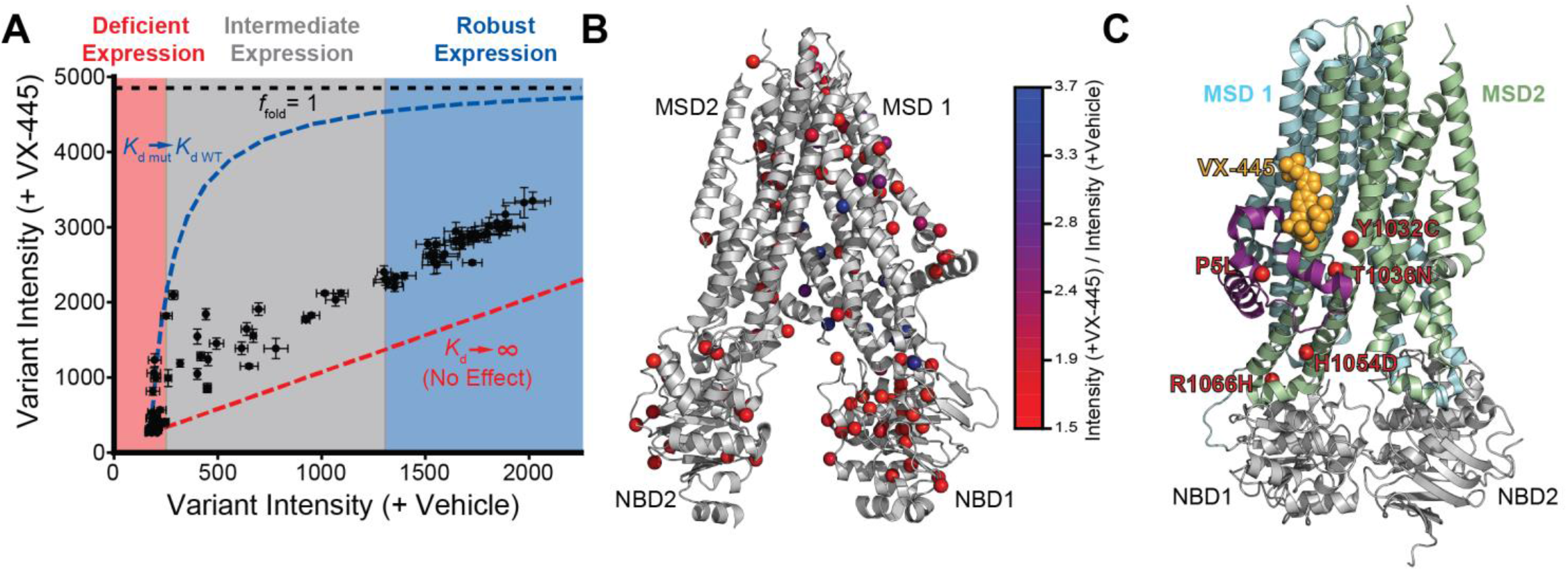
Impact of VX-445 on the Plasma Membrane Expression of CFTR Variants. Surface immunostaining intensities for individual variants were determined in the presence and absence of 3 μM VX-445 using deep mutational scanning (DMS). A) Variant immunostaining intensities in the presence of VX-445 are plotted against the corresponding intensities in the presence of vehicle. The relative positions of variants with deficient (red), intermediate (gray), and robust (blue) plasma membrane expression are shown for reference. Thermodynamic projections for the minimum (red dashes) and maximum shifts (blue dashes) in variant intensities are shown based on the projected surface immunostaining intensity of fully folded CFTR (*f*_fold_ = 1, black dashes). Measurements represent the average of three biological replicates and error bars reflect the standard deviation. B) The fold-change in intensity values for individual variants in the presence of VX-445 are projected onto their mutated side chains in the CFTR closed state structure (PDB 5UAK). Side chain C_β_/ glycine H atoms are rendered as spheres and colored according to the average fold-change from three biological replicates. C) The positions of five poorly expressed mutants that exhibit robust, selective response to VX-445 are indicated in red within the VX-445-bound CFTR active state structure (PDB 8EIG). The bound VX-445 is shown orange. The lasso domain, MSD1, and MSD2 are shown in purple, cyan, and green, respectively.

### Structural and Energetic Basis of Corrector Selectivity

To assess the structural basis of corrector selectivity, we used a recently established modeling approach to compare the effects of VX-661 binding on the structural properties of 15 CF variants that differ with respect to their sensitivity to VX-661 and VX-445 (Table S2).^26^ Briefly, we initially threaded the sequences of each variant into a set of five structural models that fit best within previously published Cryo-EM density maps of the CFTR active state.^27,28^ We then relaxed each model to generate an ensemble of low energy, native-like CF variant conformations. We next used RosettaLigand^29^ to dock VX-661 into its binding site and employed RosettaCM^30^ to generate an ensemble of 2,000 low-energy apo and bound conformations for each variant. VX-661-selective mutations such as P67L typically exhibit structural perturbations within TMD1 and within the MSD1-NBD1 interface (Fig. 6A). VX-661 binding generally enhances the formation of native structural orientations throughout the helical bundle-particularly within TMD7 and the ICL2-NBD2 interface (Fig. 6B). By comparison, the perturbations caused by certain VX-445-selective mutations such as R1066H extend further into MSD1, MSD2, and their domain interfaces (Fig. 6C). The binding of VX-661 to this variant stabilizes some of the same regions of the helical bundle observed in the P67L bound state, including TMD7, but generally fails to restore native contacts within domain interfaces (Fig. 6D). Together, these observations suggest that, while the structural effects of VX-661 binding are comparable across variants with divergent pharmacological properties, the mutations vary with respect to both the regions of the native structure they perturb and the overall number of native contacts they disrupt.

**Figure 6.**
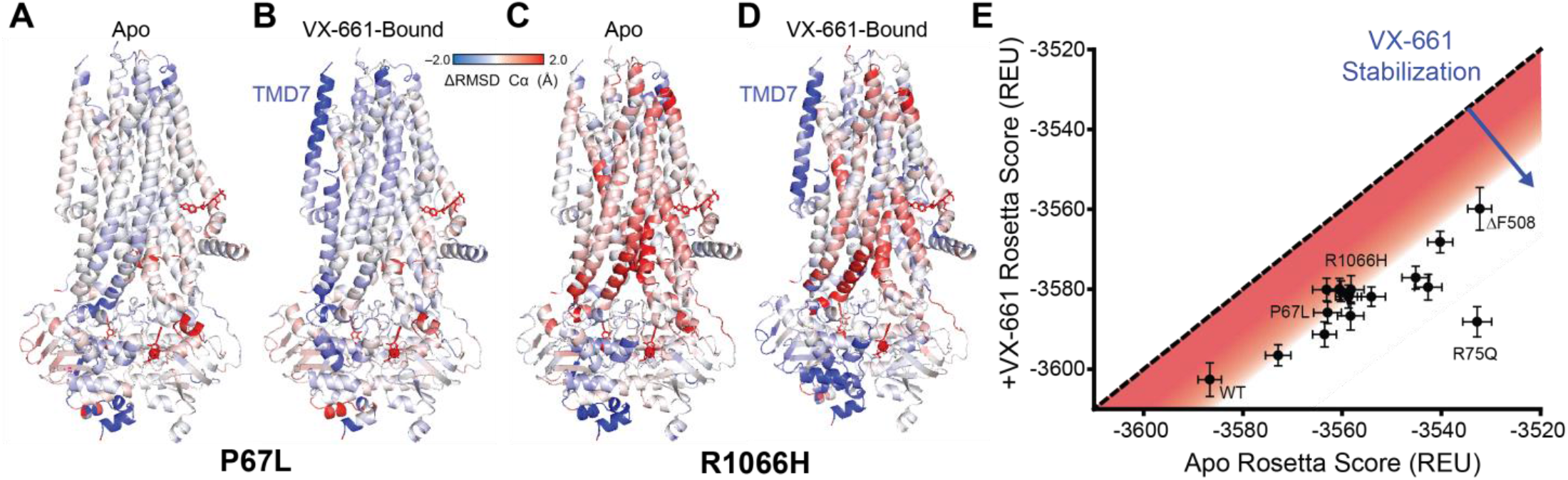
Variant-Specific Structural and Energetic Effects of VX-661 Binding. A) Low-energy structural ensembles of P67L and WT CFTR were generated and the differences in average residue Cα RMSD (mutant-WT) are projected onto PDB 7SV7. Regions where the mutant structural ensemble exhibits greater structural variation are shown in red while those that adopt an ordered orientation are shown in blue. B) An ensemble of VX-661-bound P67L CFTR models was generated and the differences in the average residue Cα RMSD (apo-bound) are projected onto PDB 7SV7. Regions where the bound mutant structural ensemble exhibits greater structural variation are shown in red while those that adopt an ordered orientation in the bound state are shown in blue. C) Low-energy structural ensembles of R1066H and WT CFTR were generated and the differences in average residue Cα RMSD (mutant-WT) are projected onto PDB 7SV7. Regions where the mutant structural ensemble exhibits greater structural variation are shown in red while those that adopt an ordered orientation are shown in blue. D) An ensemble of VX-661-bound R1066H CFTR models was generated and the differences in the average residue Cα RMSD (apo-bound) are projected onto PDB 7SV7. Regions where the bound mutant structural ensemble exhibits greater structural variation are shown in red while those that adopt an ordered orientation in the bound state are shown in blue. E) The average Rosetta energy scores of the VX-661-bound state are plotted against that of the apo CFTR variant models. A line with a slope of one intersecting the origin (no effect) is shown for reference. Changes in RMSD and the standard deviations of Rosetta energy scores were calculated across the 200 lowest energy scoring models.

VX-661 binding appears to promote the formation of a similar set of native contacts within VX-661-sensitive and insensitive variants alike (Fig. 6 B & D). Based on this consideration, we hypothesized that the magnitude of the energetic stabilization achieved by VX-661 binding is comparable across pharmacologically divergent variants. Indeed, a plot of the average Rosetta Energy Scores for the VX-661-bound variant models against the scores for their apo models suggests binding generally reduces the energy of the native conformation for variants with robust (e.g. WT) and deficient (e.g. ΔF508) PME alike (Fig. 6E). Interestingly, we did identify one mutation (R75Q) that neutralizes a charged residue adjacent to a VX-661-coordinating side chain (R74)^7^ and enhances its stabilization. Though there may be rare instances like this in which CF mutations directly perturb the binding pocket, the general trend in Fig. 6C implies selectivity is generally unlikely to emerge from specific structural features within the VX-661-bound state. Rather, the insensitivity of some variants may arise from the fact that the fixed VX-661 binding energy is insufficient to compensate for the magnitude of the destabilization caused by certain mutations (see *Discussion*). Indeed, the VX-661-insensitive variant R1066H disrupts a larger proportion of native contacts and is projected to be less stable than the VX-661-sensitive P67L CFTR (compare Figs. 6 A & C, Table S2). Nevertheless, we note that this variable alone is unlikely to explain the preference of certain variants for one corrector over another-its PME is 2.3-fold greater in the presence of VX-445 relative to that in the presence of VX-661 despite the fact that these two correctors bind with comparable affinity. The selectivity of R1066H and other variants bearing mutations within C-terminal domains for VX-445 may reflect both their degree of their destabilization and the timing of the misfolding reaction in relation to the emergence of the corrector binding site from the ribosome-translocon complex (see *Discussion*).

### Impact of Combining VX-661 and VX-445 on CFTR Variant Plasma Membrane Expression

Combining VX-661 with VX-445 restores PME to certain misfolded CFTR variants that cannot be rescued by either compound alone. To survey the combined effects of these correctors, we repeated DMS measurements in the presence of both VX-661 and VX-445. Incubating recombinant cells expressing CFTR variants in media containing 3 μM VX-661 and 3 μM VX-445 generates a substantial increase in the proportion of cells with WT-like surface immunostaining relative to either individual treatment (Fig. 1A). The average effect size of the combination treatment across the 105 potentially correctable variants in this library (2.8-fold) is much greater than that of either VX-661 (1.3-fold) or VX-445 (1.1-fold) alone (Fig. 3A). It should also be noted that this enhancement occurs across all subdomains (Fig. 2 B & D, and 7B). Interestingly, while the effects of this combination treatment are again most pronounced for variants with intermediate PME, combining these molecules also appreciably enhances the immunostaining of variants with deficient PME (Figs. 3A). Several of these poorly expressed variants (P67L, Q98R, L165S, L206W, I336K, M470V, S492S, V520F, S549N, T1036N, H1054D, R1066H, and L1077P) exhibit increases in immunostaining that are in quantitative agreement with theoretic projections based on the thermodynamic shifts in the folding equilibrium (blue dashes, Fig. 7A), which again suggests the rescue of certain variants is rooted in their intrinsic folding energetics (see *Discussion*). Overall, these 13 variants that are relatively insensitive to either molecule alone yet achieve WT immunostaining levels in the presence of both molecules (Table S1). Moreover, they are distributed throughout the structural subdomains of the CFTR protein, which implies that the total combined binding energy of these compounds is sufficient to rescue certain mutagenic defects in other subdomains (see *Discussion*).

**Figure 7.**
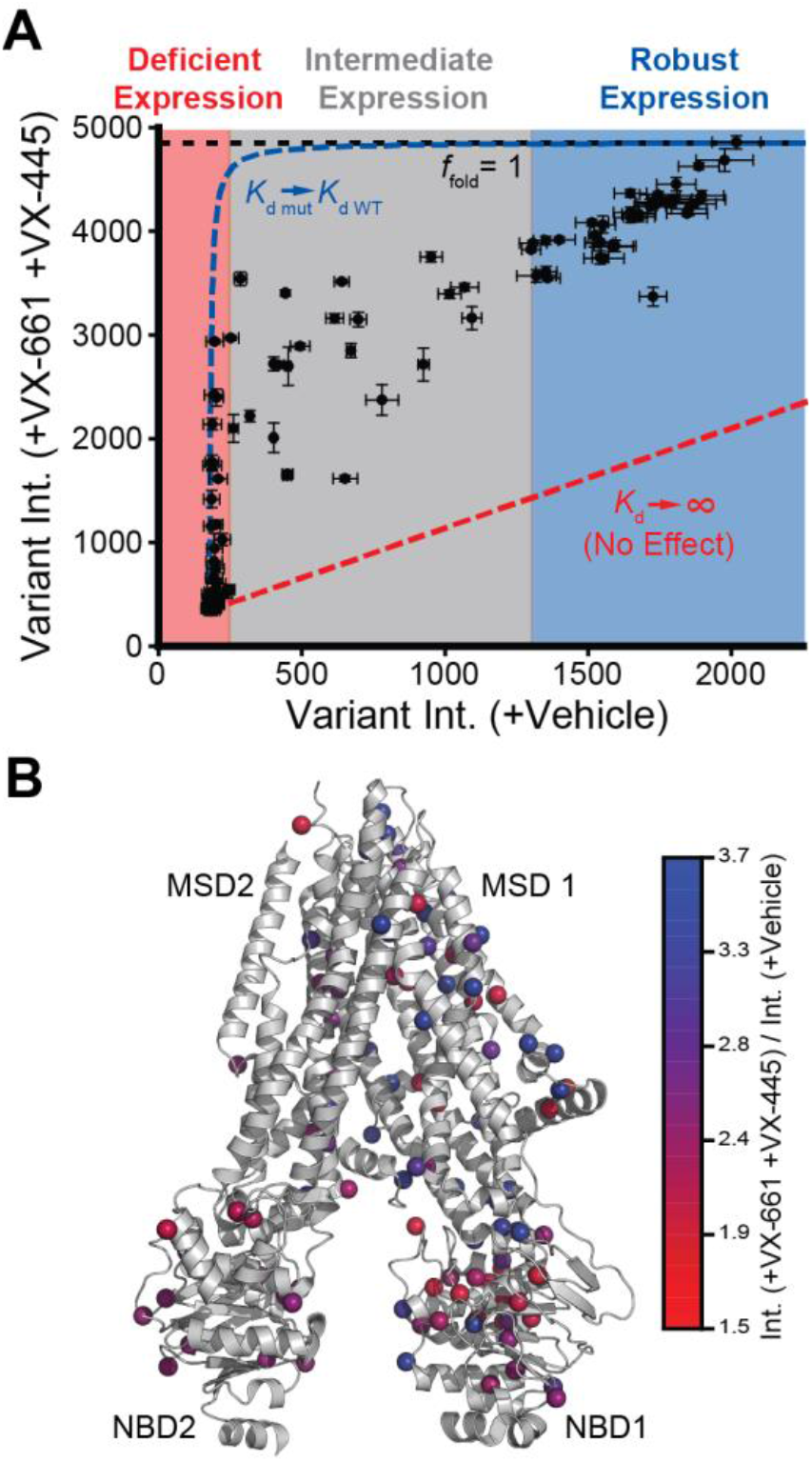
Impact of the combination of VX-661 and VX-445 on the Plasma Membrane Expression of CFTR Variants. Surface immunostaining intensities for individual variants were determined in the presence of the combination of 3 μM VX-661 and 3 μM VX-445 using deep mutational scanning (DMS). A) Variant immunostaining intensities in the presence of VX-661 + VX-445 are plotted against corresponding intensities in the presence of vehicle. The relative positions of variants with deficient (red), intermediate (gray), and robust (blue) plasma membrane expression are shown for reference. Thermodynamic projections for the minimum (red dashes) and maximum shifts (blue dashes) in variant intensities are shown based on the projected surface immunostaining intensity of fully folded CFTR (*f*_fold_ = 1, black dashes). Measurements represent the average of three biological replicates and error bars reflect the standard deviation. B) The fold-change in intensity values for individual variants in the presence of VX-445 are projected onto their mutated side chains in the CFTR closed state structure (PDB 5UAK). Side chain C_β_/ glycine H atoms are rendered as spheres and colored according to the average fold-change from three biological replicates.

## Discussion

The recent development of targeted combinations of corrector and potentiator molecules known as “highly effective modulator therapies” have revolutionized the clinical treatment of CF. Nevertheless, there are still many people with CF harboring rare CFTR variants that fail to respond to various CFTR modulators for reasons that remain unclear. Both basic insights into the mutation-specific effects of these compounds and efficient methodologies to survey the pharmacological properties of rare variants are needed to optimize and target new and improved combinations of emerging CFTR modulators. In this work, we develop a DMS approach to quantitatively compare the effects of correctors on the PME of 129 known CF variants in a single experiment. Despite the limitations associated with the use of HEK293T cells, we show that surface immunostaining profiles are broadly consistent with previous variant classifications, where available (Table S1, Fig. 1B). Across the 105 potentially correctable missense and indel variants within this library, we identify 67 variants that exhibit diminished PME (Table S1)-most of which feature mutations within MSD1 or NBD1 (Fig. 2C). VX-661 and VX-445 are particularly effective towards distinct, non-overlapping sets of variants with intermediate PME (Figs. 4B & 5B). Variants that are selective for VX-661 cluster near its binding site between TMDs 1, 2, 3, and 6 (Fig. 4C)-three segments in MSD1 that are translated during the early stages of CFTR assembly.^7^ Similarly, VX-445-selective variants cluster near its binding site within TMDs 10 & 11 (Fig. 5C)-two domain swapped segments that form their native contacts with MSD1 and NBD1 following the synthesis of the first 1,000+ CFTR residues.^13^ Combining these correctors synergistically enhances the PME of numerous CF variants bearing mutations throughout the CFTR structure (Fig. 7). We note that the heightened sensitivity of variants with intermediate PME echoes previous findings that CF variants with residual function are most susceptible to CFTR modulators.^31^ Together, these findings provide an unprecedented overview of the effects of correctors on rare CF variants. Moreover, our findings provide complementary support for structural evidence suggesting their synergistic effects arise, in part, from their stabilization of distinct structural motifs that form during different stages of CFTR synthesis.^13^

CF mutations cause a variety of conformational defects in the CFTR protein including but not limited to the perturbation of interhelical contacts and the destabilization of domain-domain interfaces.^21^ These conformational defects can occur at various points during and after its ∼5 min. cotranslational folding process in a manner that can ultimately alter the CFTR interactome and compromise its PME and/ or maturation.^24,32,33^ Correctors reverse some of these conformational defects and restore the native interactomes of certain variants through mechanisms that are potentially as diverse as the structural effects of the mutations themselves.^24^ Nevertheless, the selective binding and energetic stabilization of the native fold is generally considered to be a key ingredient for the rescue of misfolded variants by correctors and other pharmacochaperones.^34,35^ Indeed, there are several indications that binding energetics shape the selectivity of certain variants for these treatments. Most notably, we find that the effect sizes of all three treatments are generally maximized for variants with intermediate PME (Fig. 3A). This observation, which echoes our recent findings on the pharmacochaperone-mediated rescue of misfolded rhodopsin variants,^17^ implies that the ability of these compounds to re-balance the folding equilibrium and prevent off-pathway interactions is fundamentally limited, in part, by their binding energy. Our modeling efforts reveal that VX-661 binding generally enforces a common subset of native contacts and provides a similar degree of stabilization for a variety of variants with divergent properties (Fig. 6). Response to VX-661 therefore appears to have less to do with differences in the structural properties of the bound state and more to do with the nature of the misfolding reaction. Thus, while the net sensitivity of individual variants may depend on a variety of factors ranging from specific changes in the interactome to changes in corrector binding affinity, the energetic interplay between binding and folding appears to be a key consideration to maximize efficacy across the spectrum of rare variants.

The energetic interpretation of the observed trends described above has general implications for the mechanistic basis for the mutation-specific corrector effects. We approximated theoretical upper and lower bounds for their change in CF variant immunostaining based on mutant PME levels and corrector binding affinities as previously described (compare Figs. 4A, 5A, and 7A).^17^ Many variants with intermediate PME exhibit increases in intensity that approach their upper bound in the presence of VX-661 (Fig. 4A), suggesting their shifts in the folding equilibrium are effectively offset by its binding energy. However, there are a few insensitive variants that may compromise VX-661 binding and/ or trigger irreversible assembly defects. This latter possibility may explain the muted effects of VX-445 relative to VX-661 overall (compare Figs. 4A & 5A). Though these two correctors bind with comparable affinity, the stabilization of C-terminal regions by VX-445 potentially occurs minutes after the translation of MSD1 and NBD1, where most of the destabilizing mutations are located (Fig. 2C). Notably, thermodynamic limits suggest that binding energy of VX-661 or VX-445 alone is generally insufficient rescue variants with deficient PME-their maximum theoretical shift is relatively modest (Figs. 4A & 5A). However, combining these molecules generates prominent shifts in the PME of certain variants with deficient PME that are in quantitative agreement with the steeper upper boundary generated by their combined binding energy (Fig 7A). These observations suggest combinations of correctors are superior for two reasons: 1) binding at different stages of assembly may suppress kinetically decoupled assembly defects and 2) combining two molecules increases the overall binding energy beyond what can be achieved at a single pocket. Interestingly, we note that both the observed effect sizes of these compounds and the estimated upper boundaries are much greater overall for CFTR variants relative to rhodopsin variants. The stabilizing 9-*cis*-retinal cofactor increases the PME of rhodopsin variants with WT-like PME by ∼30%^36^ whereas treatment with VX-661 + VX-445 enhances CFTR variants with WT-like PME by ∼200% (Table S1). The increased “growth potential” of CF variants may reflect the intrinsic instability of the WT channel-CFTR exhibits ∼30-fold lower PME relative to rhodopsin under similar conditions when adjusted for the number of epitopes.^17^ Based on this consideration, we suspect that large, poorly expressed multi-domain proteins may generally constitute favorable targets for correctors given their relative abundance of potential binding sites and low intrinsic folding efficiencies, which render PME highly tunable.

In addition to these basic insights into the mechanistic effects of correctors, our findings provide an unprecedented overview of the pharmacological properties of rare CF variants. We note that there are four uncharacterized, “off-label” rare CF variants with deficient PME in our library that exhibit prominent increases in PME in the presence of VX-661 and VX-445 (Q359K/ T360K, P750L, L927P, and L1065P, Table S1). Follow-up investigations are needed to determine whether the enhanced PME of these variants translates to increases in CFTR function. We note that the utility of the DMS-based PME measurements described herein would be greatly enhanced by the incorporation of a complementary CFTR function assay. Unfortunately, current halide-sensitive yellow fluorescent protein (hYFP)-based conductance assays^37^ require kinetic measurements that are fundamentally incompatible with the prolonged cell sorting-based separations needed to parse the effects of these variants. Nevertheless, the assay described herein still provides an unprecedented approach to evaluate the mutation-specific effects of new combinations of CFTR modulators at scale. This DMS assay, which can be easily expanded to incorporate new emerging CF variants from diverse populations, could potentially be used to optimize the targeting of new CFTR modulators ranging from those currently in clinical trials to those that are currently under development. Furthermore, this approach could also potentially be used to re-purpose lead-compounds that were previously triaged due to their modest activity towards ΔF508. Finally, we note that this assay provides a means to build information about how these variants respond to various compounds could eventually be used to derive a knowledge-based predictive theratyping system based on the observed pharmacological properties of these variants rather than their biochemical effects.^2^ Thus, the DMS approach described herein provides an efficient tool to develop and refine precision CF therapeutics.

## Materials and Methods

### Plasmid preparation and mutagenesis

A pcDNA5 vector containing CFTR without a promoter, a triple hemagglutinin (HA) tag in the fourth extra-cellular loop, an internal ribosome entry site-eGFP cassette, and a Bxb1 recombination site was used to generate a molecular library of barcoded CF variants. We first installed a randomized ten nucleotide “barcode” region upstream of the Bxb1 recombination site using nicking mutagenesis. A plasmid preparation containing a mixed population of barcoded WT plasmid was used as a template for 129 individual site-directed mutagenesis reactions to generate a library of pathogenic CF variants that were found in the CFTR2 Database (https://www.cftr2.org/). Individual clones from each reaction were generated using the Zyppy-96 Plasmid Kit (Zymo Research, Irvine, CA) and deep sequenced to confirm the sequence of each mutated open reading frame and to determine its corresponding 10 base barcode sequence. Plasmids encoding individual variants were pooled and electroporated into electro-competent NEB10β cells (New England Biolabs, Ipswitch, MA), which were then grown in liquid culture overnight and purified using the ZymoPure endotoxin-free midiprep kit (Zymo Research, Irvine, CA). Transient expression of individual variants for western blots was carried out using a distinct set of expression vectors encoding the untagged CFTR variant cDNAs in the context of a pCDNA5 expression vector, which were generated by site-directed mutagenesis. The Bxb1 recombinase expression vector (pCAG-NLS-HA Bxb1) was kindly provided by Douglas Fowler.

### Western Blot Analysis of CFTR Expression Levels

A series of untagged CFTR variants were transiently expressed in HEK293T cells prior to the generation of cellular lysates and the analysis of expression levels by western blot. Briefly, cells were grown in 6 cm dishes in complete Dulbecco’s modified Eagle medium (Gibco, Carlsbad, CA) supplemented with 10% fetal bovine serum (Corning, Corning, NY) and penicillin (100 U/ml)/ streptomycin (100 μg/ml) (complete media) then transfected with Fugene 6 (Promega, Madison, WI). Cells were then lysed 2 days post-transfection using a lysis buffer containing 150 mM NaCl, 1% Triton X-100, and 2 mg/ mL of a protease inhibitor cocktail (Roche Diagnostics, Indianapolis, IN) in 25 mM Tris (pH 7.6). Total protein concentrations were then determined using a detergent-compatible Bradford assay (Pierce Biotechnology, Waltham, MA). Lysates were then diluted into SDS-PAGE sample buffer and heated to 37°C for 30 min prior to loading 52 μg of total protein from each sample onto a 7.5% SDS-PAGE gel. Proteins were then separated by electrophoresis and transferred onto a PVDF membrane. Membranes were blocked using a 5% milk solution in TBST buffer prior to incubating the membrane in a wash solution containing a mouse anti-CFTR antibody (1:1,000 dilution AB217, CFTR Antibody Distribution Program, Chapel Hill, NC). Membranes were then washed three times with TBST buffer prior to incubating the membrane in a wash solution containing an IRDye 680RD-labeled goat anti-mouse secondary antibody (1:10,000 dilution, LI-COR Biosciences, Lincoln, NE). The membrane was then washed three more times in TBST prior to fluorescent imaging using an Odyssey CLx system (LI-COR Biosciences, Lincoln, NE). Mature CFTR band intensities were then quantified using Image Studio V. 5.2 software (LI-COR Biosciences, Lincoln, NE). Loading controls were collected for each blot by cutting off the lower portion of the membrane and repeating the protocol using the same protocol, except a mouse anti-GAPDH primary antibody (1:10,000 dilution ab8245, Abcam, Cambridge, UK) was used in place of the anti-CFTR antibody.

### Production and fractionation of recombinant cell lines

A pool of recombinant stable cells expressing individual CF variants was generated using a previously described stable HEK293T cell line containing a genomic Tet-Bxb1-BFP “landing pad.” Recombinant cells were generated and isolated as was previously described. Briefly, cells grown in 10 cm dishes in complete media were co-transfected with our library of cystic fibrosis variants and the Bxb1 recombinase expression vectors using Fugene 6 (Promega, Madison, WI). Doxycycline (2 μg/mL) was added one day after transfection and the cells were grown at 33 °C for the following 3 days. The cells were then incubated at 37°C for 24 hours prior to the isolation of GFP positive/ BFP negative cells that had undergone recombination using a BD FACS Aria II (BD Biosciences, Franklin Lakes, NJ). These cells were grown in 10 cm dishes with complete media supplemented with doxycycline (2 μg/mL) until confluency when they were divided into 15 cm dishes for the three drug trial replicates. Where indicated, cells were incubated with either DMSO (vehicle), 3 μM VX-445, VX-661, or a combination of the two for 16 hours prior to sorting. CFTR expressed at the plasma membrane of recombinant cells was labeled with a DyLight 550–conjugated anti-HA antibody (ThermoFisher, Waltham, MA). Labelled cells were then fractionated into quartiles according to surface immunostaining intensity using a FACS Aria IIu fluorescence activated cell sorter (BD Biosciences, Franklin Lakes, NJ). At least 2 million cells from each fraction were isolated to ensure exhaustive sampling. Fractionated sub-populations were expanded in 10 cm culture dishes prior to harvesting and freezing 10-20 million cells per quartile fraction for the downstream genetic analysis.

### Extraction of genomic DNA and preparation of next-generation sequencing libraries

To track the surface immunostaining of individual CF variants, we first extracted the gDNA from each cellular fraction using the GenElute Mammalian Genomic DNA Miniprep kit (Sigma-Aldrich, St. Louis, MO). A previously described semi-nested polymerase chain reaction (PCR) technique was then used to selectively amplify the barcoded region of the recombined plasmids within the gDNA. Briefly, an initial PCR reaction was used to first amplify the region of interest from the gDNA. The product of this reaction was then used as a template for a second PCR reaction that amplified the barcoded region while installing indexed Illumina adapter sequences. Amplicons were gel-purified using the Zymoclean Gel DNA Recovery Kit (Zymo Research, Irvine, CA). The purity of each sequencing library was confirmed using an Agilent 2200 TapeStation (Agilent Technologies, Santa Clara, CA). Libraries were sequenced using a NextSeq 500 Mid Output 150-cycle kit (Illumina, San Diego, CA) at an average depth of ∼2 million reads per quartile.

### Estimation of surface immunostaining levels from deep mutational scanning data

Surface immunostaining levels were estimated from sequencing data using a computational approach described previously. Briefly, low quality reads that were likely to contain more than one error were removed from the analysis. The remaining reads containing one of the 129 barcodes corresponding to a variant of interest were then rarefied to generate subsampled datasets with a uniform number of reads for each sample. We then calculated the weighted-average immunostaining intensity values for each barcode/ variant using the following equation:

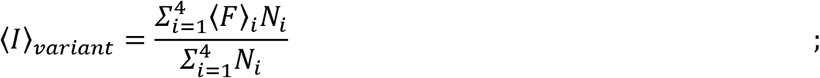

where ⟨I⟩_variant_ is the weighted-average fluorescence intensity of a given variant, ⟨F⟩_i_ is the mean fluorescence intensity associated with cells from the i^th^ FACS quartile, and N_i_ is the number of barcode/ variant reads in the i^th^ FACS quartile. Variant intensities from each replicate were normalized relative to one another using the mean surface immunostaining intensity of the recombinant cell population on each day to account for small variations in laser power and/ or detector voltage. Intensity values reported herein represent the average normalized intensity values from three replicate experiments.

### Derivation of Thermodynamic Bounds for Corrector Response

A series of simplifying assumptions was used to derive thermodynamic boundaries for the change in CFTR variant surface immunostaining intensities in the presence of correctors as previously described.^17^ Briefly, we first assumed surface immunostaining intensities to be proportional to the equilibrium fraction of folded protein (*f*_fold_) of CFTR variants. The lowest observed variant immunostaining intensity under each condition was taken as an estimate for the immunostaining intensity of cells expressing variants with an *f*_fold_ ∼ 0. The maximum overall immunostaining intensity across all datasets (G551S CFTR in the presence of VX-661 and VX-445, Int. = 4857.23) was taken as an estimate for the immunostaining intensity of cells expressing variants with an *f*_fold_ ∼ 1. These estimates were then used to approximate the equilibrium *f*_fold_ of each variant and the corresponding free energy of unfolding (ΔG_fold_). We then used the concentration of each corrector compound and their equilibrium dissociation constants (K_d, VX-661_ = 127 nM, *K*_d, VX-445_ = 67 nM, FDA New Drug Application, https://www.accessdata.fda.gov/drugsatfda_docs/nda/2019/212273Orig1s000MultidisciplineR.pdf) to estimate the effects of binding on ΔG_fold_ as well as the corresponding change in *f*_fold_ and immunostaining intensity.

### Cryo-EM Refinement, VX-661 Docking, and Template Selection

We generated an ensemble of active conformation CFTR models (PDB ID 6MSM) by refining the structure into the published cryo-EM density map with Rosetta as described previously.^26,28^ Briefly, refinement parameters were optimized in our previous study, and used to generate 2000 models. The lowest scoring five models by Rosetta score (in Rosetta energy units) were selected for molecular docking of VX-661. The ensemble of structures created during cryo-EM refinement may not contain the same coordinates as the published bound model VX-661 bound model. Hence, we docked VX-661 to each of the five lowest scoring cryo-EM refinement models 1000 times using Rosetta Ligand^29^ and took the lowest scoring model with the correct orientation in the binding pocket as a template for RosettaCM.^30^

Quantitative comparison between the apo and VX-661-bound mutant models requires simulations that are initiated from the same initial coordinates. Since VX-661 bound coordinates were not available at the time the study began, we chose to dock VX-661 to the apo templates generated above from PDB 6MSM (active apo state) instead of PDB 7SV7 (active bound state) because the 6MSM template binding pocket coordinates fail to precisely overlap with the binding pocket coordinates in 7SV7. A conformational ensemble for VX-661 was created using the BCL ConformerGenerator.^38^ BCL exported a Rosetta-readable parameter file for ligand docking with partial charges, centroid, and torsional parameter files. For docking, we positioned VX-661 at the same location as seen in the published bound model.^7^ The initial low-resolution docking was completed using the Transform mover in Rosetta with a box size of 5.0 Å, move distance of 0.1 Å, for 500 cycles. Then 1000 full atom docking simulations were performed using RosettaLigand for 6 cycles repacking every 3 cycles and interface energy scores were reported. The lowest 20 scoring interface energy models were manually inspected to determine which docked pose demonstrated overlap and correct orientation compared to the published VX-661 binding site. This model was selected for use as a final template for comparative modeling with RosettaCM.

### In Silico Mutagenesis, RosettaCM of Mutants, and Evaluation

CFTR point mutations were introduced using the MutateResidue mover in Rosetta. First, in the active state model, 6MSM, we mutated E1371 back to the naturally occurring glutamine residue. Next, mutations of interest were introduced to the templates. We generated deletion mutations by removing F508 from the active state CFTR fasta files and threading the sequence onto the active state template models. We used RosettaCM to model CFTR variants using our previously published approach.^26^ Briefly, we performed comparative modeling with static templates derived from the cryo-EM density following molecular docking of VX-661. To ensure any alterations to the structure by docking VX-661 were accounted for, we used the VX-661 docked cryo-EM refinement models as templates and removed the VX-661 atom coordinates and parameters files from the simulations of the apo state. This allowed RosettaCM to start with the same templates and only diverge by the presence of VX-661. Finally, we ran RosettaCM with multiple template hybridization using the HybridizeMover in Rosetta guided by the RosettaMembrane energy function.^39,40^ Membrane specific Rosetta energy terms were imposed within a theoretical membrane bilayer by defining the transmembrane helix regions with the prediction software OCTOPUS.^41^

We evaluated protein thermodynamic stability metrics for WT, F508del, and other CFTR variants using Rosetta score and alpha carbon (Cα) root mean squared deviation (RMSD) for whole structures as well as on a per-residue basis with respect to a reference model (either the published model or a low scoring model in the ensemble) as previously described.^26^ We calculated ΔΔG by subtracting the average Rosetta energy scoring in an apo ensemble from the average of the WT ensemble. Alternatively, we calculated the ΔΔG of VX-661 binding by subtracting the average Rosetta energy scoring in an VX-661 bound ensemble from the average of the apo ensemble.

## Acknowledgements

We thank Douglas Fowler, Jeffrey Brodsky, and Gergely Lukacs for early conceptual guidance and/ or technical input. We thank Christiane Hassel and the Indiana University Flow Cytometry Core Facility for technical support. We thank the Indiana University Center for Genomics and Bioinformatics for experimental support. This research was supported by grants from the National Institutes of Health (NIH) (R01GM129261 to J. P. S., R01GM080403 to J.M., R35GM133552 to L.P., and R00HL151965 to K. E. O.) and the Cystic Fibrosis Foundation (SCHLEB18I0 & SCHLEB19I0 to J. P. S. and OLIVER22A0-KB to K. E. O.). F. J. R. acknowledges receipt of a predoctoral fellowship from the Graduate Training Program in Quantitative and Chemical Biology at Indiana University (T32 GM109825). E. F. M. acknowledges receipt of a predoctoral fellowship from the National Heart, Lung, and Blood Institute (F31 HL162483-01A1).

