## Supplemental Materials for "Elucidation of Global Trends in the Effects of VX-661 and VX-445 on the Expression of Clinical CFTR Variants"

**Contents:**

-Figure S1
-Table S1
-Table S2
-Supplemental References

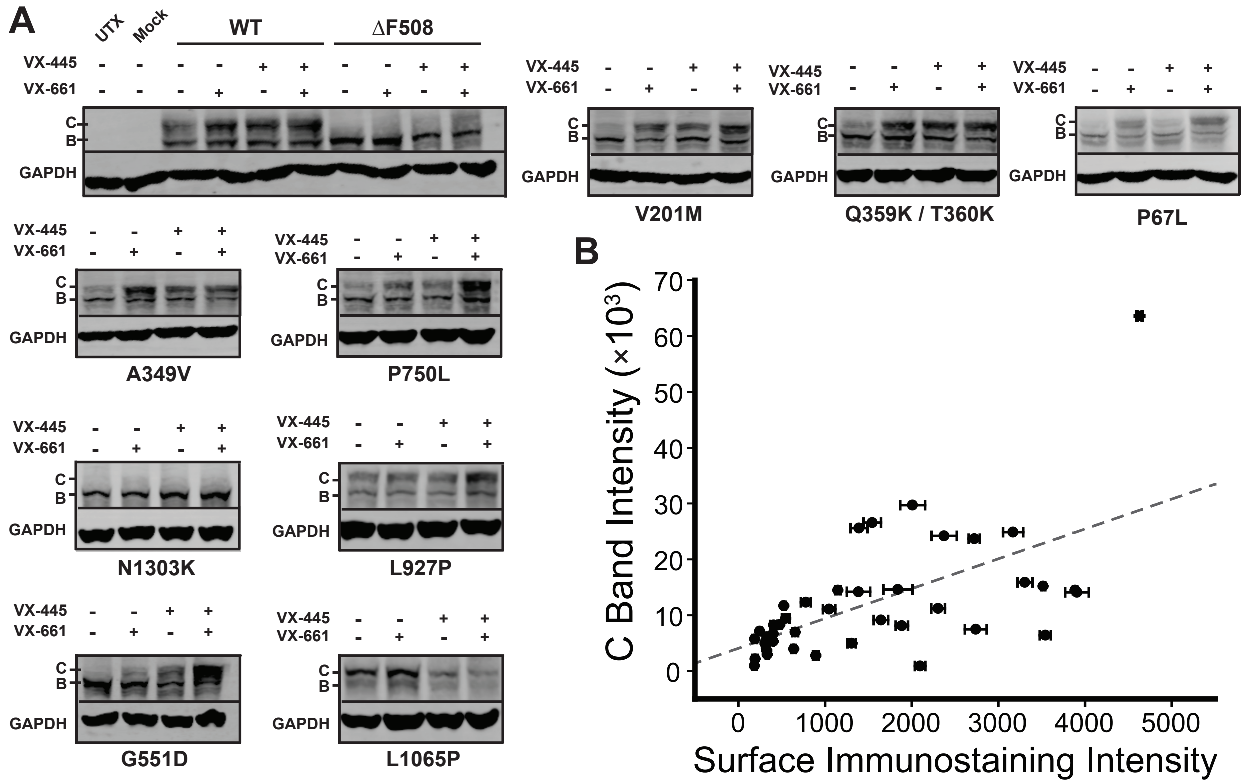


**Figure S1. Correlation Between DMS Surface Immunostaining Intensities and Mature CFTR Glycoform Levels.** A selection of ten untagged CF variants were transiently expressed in HEK293T cells prior to the analysis of CFTR expression and maturation by western blot. A) Western blots depicting the relative abundance of the immature (band B) and mature (band C) CFTR glycoforms are shown for each variant in the presence of vehicle, 3 μM VX-661, 3 μM VX-445, or 3 μM of both. A GAPDH loading control is included for reference. B) The intensity of the C band under each condition as determined by densitometry is plotted against the corresponding average surface immunostaining intensity for each variant under each condition. Error bars reflect the standard deviation from three replicates. A weighted linear fit (gray dashes) is shown for reference (Pearson's $R$ = 0.607, $p$ = 3.8 × $10^{-5}$).

### Table S1. Classifications and Surface Immunostaining Intensities of CFTR Variants

| Mutation | Putative Classification | DMS Expression | Surface Immunostaining Intensity (Apo) | Surface Immunostaining Intensity (VX-661) | Surface Immunostaining Intensity (VX-445) | Surface Immunostaining Intensity (VX-445 + VX-661) |
|---|---|---|---|---|---|---|
| P5L | Unclassified | Intermediate | 286 ± 22 | 946 ± 54 | 2097 ± 60 | 3542 ± 65 |
| R31C | Unclassified | Intermediate | 697 ± 29 | 2093 ± 103 | 1908 ± 83 | 3149 ± 71 |
| W57G | 2[2] | Deficient | 187 ± 32 | 323 ± 30 | 281 ± 36 | 368 ± 45 |
| P67L | 2, 3[1] | Deficient | 186 ± 28 | 894 ± 34 | 324 ± 44 | 1418 ± 84 |
| R74W | 2, 3[1] | Intermediate | 671 ± 15 | 2080 ± 38 | 1553 ± 88 | 2850 ± 68 |
| R75Q | Unclassified | Intermediate | 952 ± 39 | 3165 ± 95 | 1826 ± 32 | 3753 ± 49 |
| G85E | 2[1] | Deficient | 182 ± 34 | 323 ± 24 | 273 ± 33 | 364 ± 45 |
| E92K | 2, 3[1] | Deficient | 193 ± 30 | 333 ± 35 | 295 ± 33 | 420 ± 59 |
| Q98R | Unclassified | Deficient | 208 ± 32 | 1077 ± 38 | 406 ± 34 | 1613 ± 15 |
| D110H | 2, 3[1] | Robust | 1352 ± 43 | 3350 ± 82 | 2203 ± 60 | 3910 ± 40 |
| D110Y | Unclassified | Intermediate | 615 ± 30 | 2253 ± 105 | 1388 ± 82 | 3162 ± 44 |
| D110E | 3[1] | Robust | 1745 ± 65 | 3777 ± 155 | 2880 ± 78 | 4353 ± 32 |
| R117C | 2, 3[1] | Intermediate | 1068 ± 48 | 2684 ± 106 | 2032 ± 84 | 3461 ± 37 |
| R117H | 2, 3[1] | Robust | 1353 ± 36 | 3035 ± 59 | 2314 ± 70 | 3609 ± 52 |
| G126D | Unclassified | Intermediate | 453 ± 9 | 1631 ± 58 | 1243 ± 86 | 2698 ± 186 |
| I148T | Unclassified[3] | Robust | 1399 ± 56 | 3398 ± 105 | 2353 ± 42 | 3920 ± 26 |
| L165S | Unclassified | Deficient | 188 ± 32 | 352 ± 22 | 815 ± 51 | 2140 ± 55 |
| G178R | 3[1] | Robust | 1301 ± 33 | 3044 ± 141 | 2401 ± 82 | 3823 ± 24 |
| H199Y | 2[4] | Deficient | 197 ± 29 | 344 ± 32 | 306 ± 32 | 407 ± 40 |
| V201M | Unclassified | Intermediate | 639 ± 26 | 2793 ± 125 | 1645 ± 84 | 3516 ± 23 |
| P205S | 2, 3[1] | Deficient | 178 ± 28 | 341 ± 22 | 307 ± 37 | 500 ± 48 |
| L206W | 2, 3[1] | Deficient | 187 ± 29 | 624 ± 26 | 463 ± 15 | 1156 ± 25 |
| L227R | Unclassified | Deficient | 189 ± 27 | 335 ± 30 | 286 ± 40 | 386 ± 52 |
| V232D | 2, 3[1] | Deficient | 196 ± 28 | 446 ± 27 | 379 ± 42 | 943 ± 143 |
| F311L | Unclassified | Robust | 1550 ± 45 | 3220 ± 60 | 2774 ± 83 | 4065 ± 84 |
| R334W | 2, 3[1] | Robust | 1869 ± 77 | 3794 ± 141 | 2979 ± 88 | 4264 ± 39 |
| R334L | Unclassified | Deficient | 223 ± 28 | 517 ± 13 | 565 ± 32 | 1028 ± 58 |
| I336K | 2, 3[1] | Deficient | 194 ± 29 | 433 ± 22 | 365 ± 32 | 819 ± 11 |
| T338I | 2, 3[1] | Robust | 1521 ± 19 | 3291 ± 91 | 2626 ± 21 | 3958 ± 109 |
| S341P | 2, 3[1] | Intermediate | 418 ± 18 | 1572 ± 102 | 1278 ± 60 | 2711 ± 30 |
| R347H | 3[1] | Robust | 1976 ± 100 | 3902 ± 232 | 3325 ± 202 | 4686 ± 109 |
| R347P | 2, 3, 6[1] | Intermediate | 261 ± 16 | 942 ± 63 | 991 ± 114 | 2099 ± 132 |
| A349V | Unclassified[†] | Robust | 1307 ± 48 | 3306 ± 86 | 2304 ± 79 | 3884 ± 25 |
| R352Q | 3[1] | Robust | 1845 ± 71 | 3566 ± 102 | 3056 ± 109 | 4170 ± 23 |
| Q359K / T360K | Unclassified | Intermediate | 402 ± 4 | 1392 ± 99 | 1544 ± 101 | 2722 ± 64 |
| A455E | 2, 3[1] | Deficient | 192 ± 30 | 350 ± 19 | 308 ± 30 | 474 ± 62 |
| V456A | Unclassified | Deficient | 190 ± 31 | 326 ± 23 | 299 ± 28 | 480 ± 57 |
| L467P | 2[1] | Deficient | 199 ± 22 | 342 ± 17 | 292 ± 28 | 407 ± 45 |
| M470V | Unclassified | Robust | 1726 ± 48 | 3232 ± 34 | 2526 ± 29 | 3370 ± 91 |
| S492F | 2, 6[1] | Deficient | 190 ± 24 | 330 ± 25 | 291 ± 40 | 444 ± 35 |

| Mutation | Putative Classification | DMS Expression | Surface Immunostaining Intensity (Apo) | Surface Immunostaining Intensity (VX-661) | Surface Immunostaining Intensity (VX-445) | Surface Immunostaining Intensity (VX-445 + VX-661) |
|---|---|---|---|---|---|---|
| I502T | Unclassified | Deficient | 185 ± 27 | 319 ± 30 | 279 ± 28 | 426 ± 50 |
| V520F | 2[1] | Deficient | 188 ± 28 | 327 ± 30 | 281 ± 37 | 377 ± 53 |
| S549N | 3[1] | Robust | 1739 ± 69 | 3641 ± 124 | 2916 ± 88 | 4264 ± 24 |
| S549R* | 2, 3[1] | Intermediate | 450 ± 19 | 1080 ± 85 | 860 ± 61 | 1660 ± 31 |
| G551S | 3[1] | Robust | 2018 ± 85 | 4142 ± 145 | 3351 ± 114 | 4857 ± 65 |
| G551D | 3[1] | Robust | 1886 ± 71 | 3907 ± 136 | 3169 ± 116 | 4627 ± 30 |
| L558S | Unclassified | Deficient | 180 ± 24 | 310 ± 20 | 264 ± 33 | 356 ± 58 |
| A559T | 2[1] | Deficient | 179 ± 29 | 326 ± 26 | 268 ± 29 | 380 ± 50 |
| R560K | Unclassified | Deficient | 181 ± 29 | 315 ± 25 | 274 ± 28 | 379 ± 44 |
| R560T | 2[1] | Deficient | 182 ± 25 | 323 ± 25 | 281 ± 35 | 372 ± 56 |
| A561E | 2, 3, 6[1] | Deficient | 199 ± 30 | 348 ± 26 | 300 ± 32 | 401 ± 54 |
| V562I | Unclassified[†] | Robust | 1319 ± 69 | 2811 ± 70 | 2263 ± 76 | 3570 ± 58 |
| Y563N | Unclassified | Deficient | 187 ± 22 | 326 ± 26 | 277 ± 29 | 375 ± 54 |
| Y569D | 2[1] | Deficient | 185 ± 30 | 318 ± 27 | 281 ± 37 | 384 ± 47 |
| P574H | Unclassified | Deficient | 198 ± 30 | 372 ± 18 | 357 ± 29 | 748 ± 36 |
| G576A | Unclassified[†] | Robust | 1852 ± 62 | 3699 ± 148 | 2995 ± 83 | 4217 ± 63 |
| D579G | 2, 3[1] | Intermediate | 449 ± 14 | 1119 ± 71 | 852 ± 48 | 1644 ± 11 |
| H609R | 2[3] | Deficient | 185 ± 28 | 332 ± 18 | 289 ± 33 | 432 ± 44 |
| D614G | 2[1] | Deficient | 202 ± 32 | 393 ± 35 | 340 ± 35 | 604 ± 35 |
| G628R | Unclassified | Deficient | 193 ± 30 | 356 ± 31 | 316 ± 24 | 483 ± 51 |
| R668C | 2, 3[1] | Robust | 1537 ± 49 | 3218 ± 111 | 2588 ± 93 | 3884 ± 11 |
| P750L | Unclassified | Intermediate | 402 ± 7 | 1149 ± 30 | 1046 ± 71 | 2010 ± 144 |
| V754M | Unclassified | Robust | 1784 ± 72 | 3593 ± 126 | 2895 ± 75 | 4280 ± 53 |
| L927P | 2, 3[1] | Intermediate | 780 ± 57 | 1841 ± 167 | 1385 ± 134 | 2374 ± 147 |
| S945L | 2, 3[1] | Intermediate | 319 ± 6 | 981 ± 101 | 1187 ± 49 | 2218 ± 55 |
| L967S | Unclassified | Intermediate | 924 ± 20 | 2064 ± 26 | 1767 ± 32 | 2716 ± 159 |
| G970R | Unclassified | Robust | 1890 ± 90 | 3739 ± 162 | 2994 ± 103 | 4303 ± 40 |
| G970D | Unclassified | Robust | 1555 ± 70 | 3140 ± 150 | 2496 ± 111 | 3737 ± 32 |
| S977F | 2, 3[1] | Robust | 1513 ± 57 | 3205 ± 83 | 2774 ± 64 | 4083 ± 29 |
| L997F | 2, 3[1] | Robust | 1647 ± 57 | 3467 ± 145 | 2945 ± 121 | 4366 ± 37 |
| I1027T | Unclassified | Robust | 1358 ± 44 | 2947 ± 71 | 2339 ± 2 | 3548 ± 23 |
| Y1032C | Unclassified | Intermediate | 252 ± 27 | 772 ± 40 | 1818 ± 36 | 2972 ± 20 |
| T1036N | Unclassified | Deficient | 192 ± 32 | 430 ± 24 | 1057 ± 70 | 2423 ± 22 |
| F1052V | 3[1] | Robust | 1677 ± 59 | 3507 ± 124 | 2855 ± 143 | 4164 ± 74 |
| H1054D | 2, 3[1] | Deficient | 196 ± 30 | 481 ± 24 | 1230 ± 86 | 2936 ± 22 |
| G1061R | Unclassified | Deficient | 201 ± 24 | 360 ± 36 | 454 ± 20 | 1175 ± 39 |
| L1065P | 2[1] | Deficient | 194 ± 30 | 335 ± 19 | 329 ± 31 | 656 ± 34 |
| R1066C | 2[1] | Deficient | 185 ± 24 | 317 ± 28 | 283 ± 30 | 397 ± 49 |
| R1066H | 2, 3[1] | Deficient | 202 ± 25 | 439 ± 31 | 989 ± 48 | 2396 ± 82 |
| G1069R | Unclassified[†] | Robust | 1677 ± 55 | 3476 ± 118 | 2768 ± 65 | 4141 ± 23 |
| R1070W | 2, 3[1] | Intermediate | 494 ± 34 | 1588 ± 128 | 1451 ± 75 | 2891 ± 29 |
| R1070Q | 2, 3[1] | Robust | 1896 ± 76 | 3750 ± 102 | 3012 ± 65 | 4344 ± 50 |

| Mutation | Putative Classification | DMS Expression | Surface Immunostaining Intensity (Apo) | Surface Immunostaining Intensity (VX-661) | Surface Immunostaining Intensity (VX-445) | Surface Immunostaining Intensity (VX-445 + VX-661) |
|---|---|---|---|---|---|---|
| F1074L | 2, 3[1] | Intermediate | 443 ± 5 | 1710 ± 98 | 1843 ± 74 | 3405 ± 30 |
| L1077P | 2, 3, 6[1] | Deficient | 187 ± 24 | 345 ± 29 | 536 ± 24 | 1774 ± 69 |
| W1098R | Unclassified | Deficient | 201 ± 29 | 400 ± 25 | 301 ± 38 | 410 ± 45 |
| M1101K | 2[1] | Deficient | 205 ± 24 | 377 ± 26 | 314 ± 42 | 474 ± 34 |
| D1152H | 3[1] | Robust | 1542 ± 28 | 3243 ± 123 | 2536 ± 163 | 3747 ± 64 |
| L1156F* | Unclassified | Robust | 1647 ± 59 | 3454 ± 109 | 2802 ± 103 | 4183 ± 50 |
| S1159F | Unclassified | Intermediate | 1016 ± 40 | 2483 ± 106 | 2120 ± 26 | 3395 ± 41 |
| R1162L | Unclassified | Intermediate | 1093 ± 35 | 2519 ± 73 | 2119 ± 44 | 3163 ± 113 |
| I1234V | Unclassified | Robust | 1716 ± 55 | 3500 ± 127 | 2870 ± 81 | 4242 ± 68 |
| S1235R | 3[1] | Robust | 1755 ± 46 | 3597 ± 90 | 2884 ± 66 | 4293 ± 24 |
| G1244E | 3[1] | Robust | 1807 ± 68 | 3738 ± 114 | 3015 ± 114 | 4455 ± 65 |
| T1246I | Unclassified | Robust | 1724 ± 62 | 3626 ± 127 | 2902 ± 88 | 4280 ± 77 |
| S1251N | 3[1] | Robust | 1800 ± 61 | 3680 ± 140 | 2914 ± 115 | 4312 ± 43 |
| S1255P | 3[1] | Robust | 1594 ± 72 | 3331 ± 97 | 2643 ± 114 | 3859 ± 54 |
| I1269N | Unclassified | Intermediate | 651 ± 42 | 1459 ± 77 | 1148 ± 28 | 1615 ± 28 |
| D1270N | 2, 3[1] | Robust | 1585 ± 72 | 3314 ± 170 | 2609 ± 61 | 3851 ± 45 |
| Q1291H | Unclassified | Robust | 1648 ± 58 | 3450 ± 83 | 2826 ± 103 | 4130 ± 46 |
| N1303K | 2, 3, 6[1] | Deficient | 246 ± 19 | 478 ± 23 | 408 ± 41 | 547 ± 44 |
| L1335P | Unclassified | Deficient | 235 ± 22 | 428 ± 23 | 381 ± 36 | 509 ± 49 |
| G1349D | 3[1] | Robust | 1543 ± 53 | 3277 ± 24 | 2640 ± 86 | 3893 ± 114 |
| L138ins | Unclassified | Deficient | 187 ± 33 | 550 ± 11 | 453 ± 46 | 1741 ± 51 |
| I507del | Unclassified | Deficient | 186 ± 26 | 311 ± 17 | 267 ± 32 | 347 ± 49 |
| F508del | 2, 3, 6[1] | Deficient | 188 ± 28 | 331 ± 23 | 308 ± 34 | 525 ± 19 |
| T854= | Synonymous | Robust | 1721 ± 62 | 3512 ± 108 | 2840 ± 102 | 4165 ± 35 |
| Q996= | Synonymous | Robust | 1695 ± 83 | 3454 ± 100 | 2841 ± 89 | 4178 ± 22 |
| L1156= | Synonymous | Robust | 1721 ± 56 | 3554 ± 106 | 2801 ± 64 | 4129 ± 50 |
| E60X | 1 | Deficient | 184 ± 30 | 324 ± 23 | 270 ± 33 | 354 ± 50 |
| Q493X | 1 | Deficient | 191 ± 29 | 320 ± 22 | 284 ± 30 | 375 ± 62 |
| G542X | 1 | Deficient | 188 ± 32 | 315 ± 22 | 278 ± 26 | 366 ± 57 |
| R553X | 1 | Deficient | 191 ± 27 | 336 ± 27 | 285 ± 25 | 375 ± 42 |
| S912X | 1 | Deficient | 183 ± 30 | 317 ± 27 | 276 ± 24 | 359 ± 57 |
| W1089X | 1 | Deficient | 190 ± 29 | 349 ± 28 | 298 ± 24 | 373 ± 43 |
| Y1092X* | 1 | Deficient | 193 ± 34 | 337 ± 28 | 292 ± 33 | 392 ± 56 |
| E1104X | 1 | Deficient | 186 ± 25 | 315 ± 26 | 284 ± 32 | 382 ± 47 |
| R1158X | 1 | Deficient | 212 ± 30 | 386 ± 30 | 334 ± 34 | 464 ± 45 |
| R1162X | 1 | Deficient | 220 ± 28 | 389 ± 38 | 326 ± 47 | 445 ± 39 |
| S1196X | 1 | Robust | 1342 ± 22 | 2506 ± 93 | 2103 ± 125 | 2775 ± 254 |
| W1204X* | 1 | Robust | 1203 ± 26 | 2239 ± 117 | 1756 ± 153 | 2264 ± 297 |
| W1282X | 1, 2, 3, 6[1] | Deficient | 382 ± 15 | 880 ± 18 | 745 ± 83 | 1069 ± 137 |
| Q1313X | 1 | Deficient | 197 ± 30 | 354 ± 49 | 297 ± 37 | 410 ± 50 |

* These values represent the average number for multiple genetic variants that generate the same substitution.
† These variants are unclassified but were previously shown to retain the characteristics of WT CFTR.

### Table S2. Rosetta Scores of Apo and VX-661-Bound CFTR Variants

| Variant | Surface Immunostaining (DMS) | Expression Classification | Rosetta Score Apo (Rosetta Energy Units) | Rosetta Score VX-661-Bound (Rosetta Energy Units) |
|---|---|---|---|---|
| WT | 1695 ± 291* | Robust | -3587 ± 33 | -3603 ± 59 |
| P5L | 286 ± 22 | Intermediate | -3560 ± 36 | -3580 ± 36 |
| P67L | 186 ± 28 | Deficient | -3563 ± 40 | -3586 ± 38 |
| R75Q | 952 ± 39 | Intermediate | -3533 ± 41 | -3588 ± 53 |
| Q98R | 208 ± 32 | Deficient | -3563 ± 40 | -3580 ± 40 |
| L165S | 188 ± 32 | Deficient | -3561 ± 34 | -3580 ± 38 |
| V201M | 639 ± 26 | Intermediate | -3543 ± 39 | -3579 ± 46 |
| L206W | 187 ± 29 | Deficient | -3558 ± 38 | -3587 ± 49 |
| V232D | 196 ± 28 | Deficient | -3545 ± 39 | -3577 ± 40 |
| I336K | 194 ± 29 | Deficient | -3564 ± 34 | -3591 ± 45 |
| F508del | 188 ± 28 | Deficient | -3532 ± 34 | -3560 ± 76 |
| Y1032C | 252 ± 27 | Intermediate | -3573 ± 36 | -3597 ± 37 |
| T1036N | 192 ± 32 | Deficient | -3559 ± 37 | -3582 ± 41 |
| H1054D | 196 ± 30 | Deficient | -3554 ± 40 | -3582 ± 34 |
| R1066H | 202 ± 25 | Deficient | -3558 ± 37 | -3580 ± 50 |
| L1077P | 187 ± 24 | Deficient | -3540 ± 36 | -3568 ± 38 |

*Measured independently in stable cell lines expressing WT CFTR for the sake of comparison.
